# A transgenic zebrafish model for the *in vivo* study of the blood and choroid plexus brain barriers using claudin 5

**DOI:** 10.1101/180653

**Authors:** Lisanne Martine van Leeuwen, Robert J. Evans, Kin Ki Jim, Theo Verboom, Xiaoming Fang, Aleksandra Bojarczuk, Jarema Malicki, Simon Andrew Johnston, Astrid Marijke van der Sar

## Abstract

The central nervous system (CNS) has specific barriers that protect the brain from potential threats and tightly regulate molecular transport. Despite the critical functions of the CNS barriers, the mechanisms underlying their development and function are not well understood, and there are very limited experimental models for their study. Claudin 5 is a tight junction protein required for blood brain barrier (BBB) and choroid plexus (CP) barrier structure and function in humans. Here, we show that the gene *claudin 5a* is the zebrafish orthologue with high fidelity expression, in the BBB and CP barriers, that demonstrates the conservation of the BBB and CP between humans and zebrafish. Expression of *claudin 5a* correlates with developmental tightening of the BBB and is restricted to a subset of the brain vasculature clearly delineating the BBB. We show that *claudin 5a* expressing cells of the CP are ciliated ependymal cells that drive fluid flow in the brain ventricles. Finally, we find that CP development precedes BBB development and that *claudin 5a* expression occurs simultaneously with angiogenesis. Thus, our novel transgenic zebrafish represents an ideal model to study CNS barrier development and function, critical in understanding the mechanisms underlying CNS barrier function in health and disease.

## INTRODUCTION

The central nervous system is protected by three specialized barriers that shield the vulnerable brain tissue from potential threats and actively regulate exchange of ions and nutrients. The blood brain barrier (BBB) is formed by endothelial cells between blood and brain interstitial fluid and has extensive control over the immediate microenvironment of the CNS (Abbott et al., 2010, 2006). Less studied are the blood-cerebrospinal fluid (CSF) barrier, formed by the epithelial cell layer of the choroid plexus (CP) between blood and ventricular CSF, and the epithelial cell layer of the meningeal arachnoid between blood and subarachnoid CSF (Abbott et al., 2010, 2006; Obermeier et al., 2013).

The BBB and blood-CSF barrier tissues have tight junctions (TJs) consisting of protein complexes that seal adjacent cells and actively regulate barrier integrity (Greene and Campbell, 2016). TJs are protein complexes containing occludins and claudins that provide a physical barrier to block free paracellular diffusion of solutes and macromolecules (Abbott et al., 2010). More than 20 different claudin isoforms are known of which at least four, Claudin 1, 3, 5 and 12, are involved in establishing and regulating TJs in mammalian brain endothelial cells (Abdelilah-Seyfried, 2010; Greene and Campbell, 2016). Of these, *claudin 5* is the most strongly expressed in mammalian brain microvessels (Zhang et al., 2012). Whereas this protein was shown to be important for barrier integrity in mice, the expression of *claudin 5* is not conserved in the murine choroid plexus and is not a definitive marker of the BBB (Nitta et al., 2003).

Links with BBB breakdown or dysfunction have been shown for neurodegenerative processes, inflammation and infection. These include, but are not limited to, Alzheimer’s disease (Schenk and de Vries, 2016; Zenaro et al., 2016), multiple sclerosis (Macrez et al., 2016; Schenk and de Vries, 2016), amyotrophic lateral sclerosis (Garbuzova-Davis et al., 2012; Garbuzova-Davis and Sanberg, 2014); vascular dementia (Ueno et al., 2016), autoimmune encephalitis (Platt et al., 2017), and infectious meningoencephalitis (Coureuil et al., 2017; Gibson and Johnston, 2015; Swanson and McGavern, 2015). Targeting the breakdown or dysfunction of the BBB in CNS disease has significant potential as treatment target, but is severely hampered by a lack of experimental models and current treatment is limited to broad spectrum immunosuppression/anti-inflammatory treatment with, for example, glucocorticosteroids (Obermeier et al., 2013). Furthermore, to improve drug delivery in CNS disease specific modulation of therapeutic delivery to the brain with limited neurotoxicity is critical (Greene and Campbell, 2016), but there are very limited experimental models that allow the required non-invasive imaging and mechanistic studies (Greene and Campbell, 2016; O’Keeffe and Campbell, 2016; Zhang et al., 2010).

*In vitro* experimental models exist to study different aspects of the BBB, but none of them can completely mimic the complex interplay between BBB and other cells, such as immune cells or pathogens. In addition, *in vivo* systems are often restricted by the use of a single method, like single molecular tracer injections or immunohistochemistry. Moreover, real time *in vivo* imaging of the BBB and CP is not possible in current models (Blanchette and Daneman, 2015). Therefore, we set out to develop and use a new model with the potential to understand the mechanistic biology of the BBB and CP.

The zebrafish (*Danio rerio*) model has proven to be highly accessible for real time imaging especially in combination with fluorescently labelled tissues and is therefore extensively used to study development and many aspects of human disease (Holtzman et al., 2016). Zebrafish possess a BBB and CP, similar to higher vertebrates, and Claudin 5a has been suggested to play an essential role in establishment and maintenance of these barriers between systemic circulation and CNS (Henson et al., 2014; Jeong et al., 2008; Xie et al., 2010). Early in development, at 14 hours post fertilization (hpf), *claudin 5a* is expressed in the entire developing zebrafish CNS (Zhang et al., 2010). However soon after, labelling is confined to the CP and brain vasculature (from 20 hpf respectively 48 hpf onwards) (Xie et al., 2010; Zhang et al., 2010). Functional studies have shown size-dependent exclusion of fluorescent tracers injected in the circulation from 2 dpf onwards indicative for the functional maturation of the BBB shortly after TJ formation (Fleming et al., 2013; Jeong et al., 2008; van Leeuwen et al., 2014; Xie et al., 2010). Furthermore, Claudin 5a has been suggested to be involved in the establishment of the neuroepithelial ventricular barrier, which is essential for brain ventricle expansion and subsequent brain development (Zhang et al., 2012, 2010). With the use of several enhancer trap lines the presence of a diencephalic and myelencephalic CP has been suggested to be present in zebrafish larvae early in development (Bill and Korzh, 2014; Bill et al., 2008; García-Lecea et al., 2008; Henson et al., 2014). Therefore, we considered Claudin 5a to be an excellent candidate as the basis for our new *in vivo* model for the BBB and CP.

In this study, we have identified and confirmed *claudin 5a* as the zebrafish gene equivalent to human *claudin 5*. We have generated a *claudin 5a* reporter transgenic that has high fidelity of expression in both the BBB and CP. *Claudin 5a* expressing cells in the CP are ciliated ependymal cells that drive fluid flow early in development. Using our new transgenic we demonstrate that the CP forms prior to expression of *claudin 5a* in brain blood vessels but that once the BBB is established *claudin 5a* expression coincides with new BBB vessel formation.

## RESULTS

### Zebrafish *claudin 5a* is the human *claudin 5* orthologue

In order to identify the zebrafish (Dr) orthologue of human (Hs) protein Claudin 5 (*cldn5* gene) we used a BLASTP search of the zebrafish genome (GRCz10) using the protein sequence of Hs Claudin 5. Two zebrafish proteins, Claudin 5a and Claudin 5b, were identified as being most similar by protein sequence (56.9% and 54.8% identical respectively; Figure 1A). Alignment of the zebrafish protein sequences to the Hs sequence could not identify which of these two proteins is the orthologue. Examination of the genomic region and closer gene relatives of *claudin 5a* and *5b* clearly showed that *claudin 5a* shared synteny with Hs *cldn5* and that *claudin 5b* was only present in ray-finned fish (Figure 1B and C; data not shown). In addition, examination of zebrafish expression patterns (Thisse et al., 2004) showed that *claudin 5a* was expressed in the central nervous system ventricle region while *claudin 5b* had a cardiovascular patterning. Together, we took this as sufficient evidence that *claudin 5a* was the correct target gene and using BAC recombineering (Abe et al., 2011) we inserted EGFP at the translation start site of the *claudin 5a* gene with approximately 200 Kb of flanking sequence to maximise fidelity of EGFP expression to endogenous *claudin 5a* (Figure S1).

**Figure 1:**
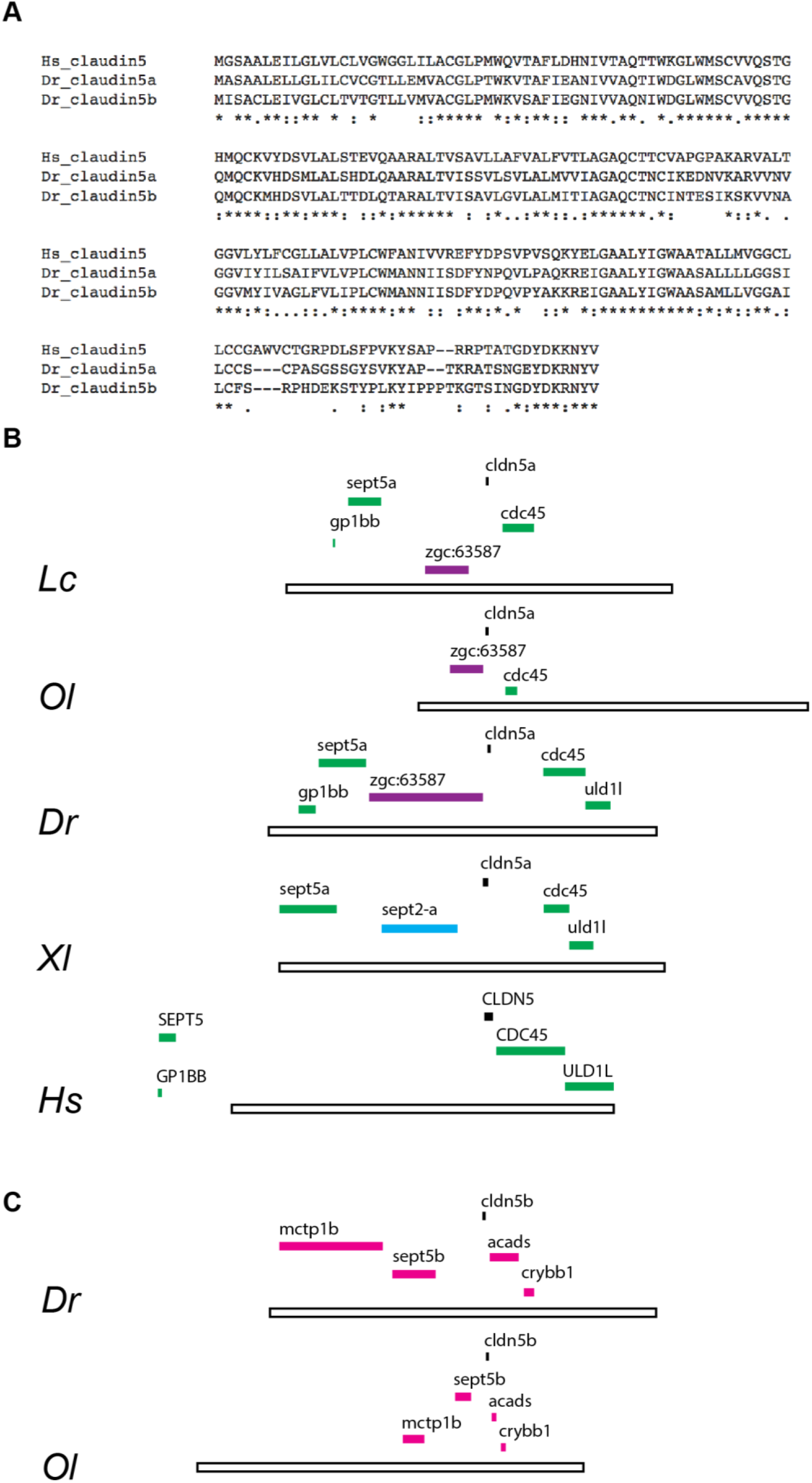
Zebrafish claudin5a is the homologue of human claudin5. A. Protein sequence alignment of human (Hs; *Homo sapiens*) claudin 5, zebrafish (Dr; *Danio rerio*) claudin5a and claudin5b using Clustal Omega. B, C. Syntenic analysis of claudin5a (B) and claudin5b (C) genes. B. Green: Syntenic genes in the claudin5a locus across *Latimeria chalumnae* (Lc), *Oryzias latipes* (Ol), *Danio rerio* (Dr), *Xenopus laevis* (Xl) and Homo sapiens (Hs). Purple: Syntenic genes in the claudin5a locus across Osteichthyes (Lc, Ol, Dr). Black bar 250 Kbases. C. Claudin5b is only present in actinopterygii (ray-finned fish), here represented by *Oryzias latipes* (Ol) and *Danio rerio* (Dr), which show conserved synteny.

### *Claudin 5a* is expressed in the choroid plexus of zebrafish at 1 dpf

To study the developmental expression of *claudin 5a* in *TgBAC(cldn5a:EGFP)*^*vum1*^ larvae and correlate this to previous performed immunohistochemical analysis (Xie et al., 2010; Zhang et al., 2012), we performed non-invasive imaging of the brain region of larvae daily between 1 and 9 days post fertilisation (dpf; Figure 2). As early as 24 hpf, GFP expression was observed in the area of the myelencephalic choroid plexus (mCP) and diencephalic CP (dCP) (Figure 2B, arrows). The mCP consisted of a large sheet of cells covering the roof of the hindbrain ventricle that developed into a compact cluster located in the midline of the larval head. In addition to expression in both CPs, labelling in brain parenchyma, presumably co-localizing with vasculature, and spinal cord was observed from 3 dpf onwards (Figure 2D, arrow). Between 3 and 5 dpf *claudin 5a* expression rapidly expanded in the entire parenchyma and had covered the complete CNS vasculature by 5 dpf (Fig 2F). Interestingly, strong labelling in the midline of the larval head was observed (Figure 2B, open arrow). This labelling appeared at the same time as in both CPs, connecting these structures, but did not co-localize with blood vessels and was sustained through development (Figure 2J, J’). Although Claudin 5a specifically labels CNS barriers in zebrafish, transient expression was observed in the caudal hematopoietic tissue (CHT), the tip of the tail and the heart region (Supplemental Figure S2). This expression was only present during early development and disappeared in later larval stages (data not shown). Collectively, our *TgBAC(Cldn5a:EGFP)*^*vum1*^ zebrafish had specific expression in brain vasculature and both CPs labelling the BBB and blood-CSF barrier respectively.

**Figure 2:**
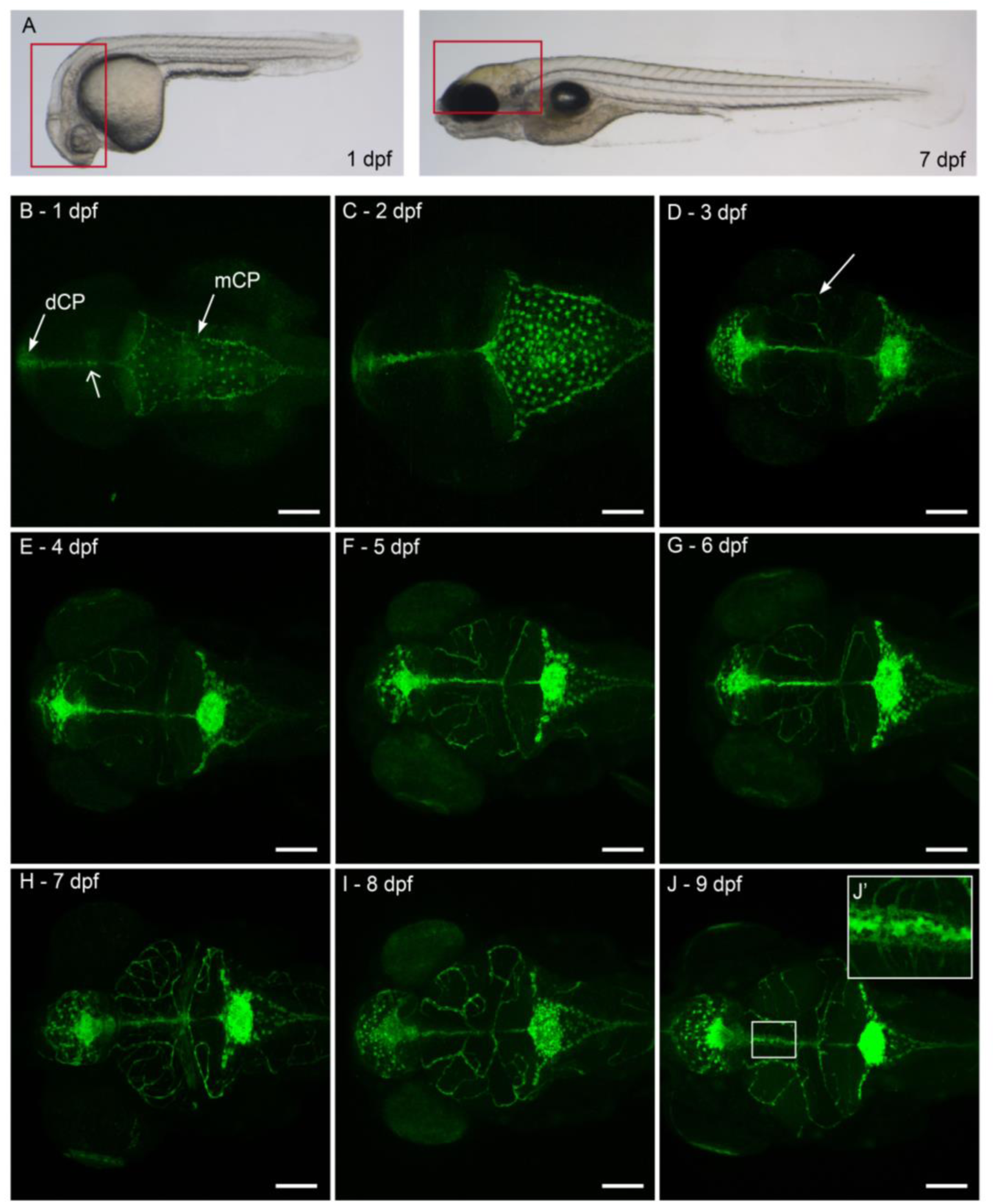
Developmental expression of claudin5a. A. Lateral view of a Casper zebrafish larva at 1 dpf and 7 dpf. Boxed area represents the brain region of which confocal images are shown in B-J. B-J. Z-stacks of dorsal view of larval head to visualize development of GFP expression from 1 to 9 dpf. GFP expression can be found in the diencephalic and myelencephalic choroid plexus (dCP, mCP) from 1 dpf onwards (B, closed arrows). In addition, labelling is observed in the midline connecting the dCP and mCP (B, arrowhead, J’). From 3 dpf onwards, labelling in brain parenchyma is observed (D, arrow). Scale Bar C-J = 100 μm

### Claudin 5a expression in brain vasculature rapidly expands between 3 dpf and 4 dpf

To study if Claudin 5a can be found in tight junctions of brain vasculature and therefore represents the BBB, we injected our construct in the vascular specific reporter line *Tg(kdrl:mCherry)*^*is5*^ (Jin, 2005) to generate a double transgenic line *Tg(kdrl:mCherry)*^*is5*^*;TgBAC(cldn5a:EGFP)*^*vum2*^. Using the previously described detailed anatomical description of vasculature development (Isogai et al., 2001) we observed that expression of *claudin 5a* first appeared in the mesencephalic vein (MsV) and middle cerebral vein (MCeV) at 3 dpf (Figure 3 A, B). Subsequent expansion of *claudin 5a* expression between 3 and 4 dpf occurred in large vessels first (Figure 3 C, D). At 5 dpf, nearly all vessels, veins and arteries, show green fluorescence indicating that *claudin 5a* is expressed in virtually all the vessels in the zebrafish brain (Figure 3 E, F). Intriguingly, a certain number of specific areas never showed *claudin 5a* expression (15 out of 16 larvae, 3 biological independent experiments, Figure 3G): the primordial midbrain channels (PMBC), choroidal vascular plexus (CVP) (Figure 3G, arrows) and anterior cerebral vein frontally located (AceV, Figure 3I-K), and at the location of the midbrain the dorsal midline junction (DMJ) and dorsal longitudinal vein (DLV) (Figure 3L-N). These Claudin 5a deficient regions were sustained through development until at least 9 dpf.

**Figure 3:**
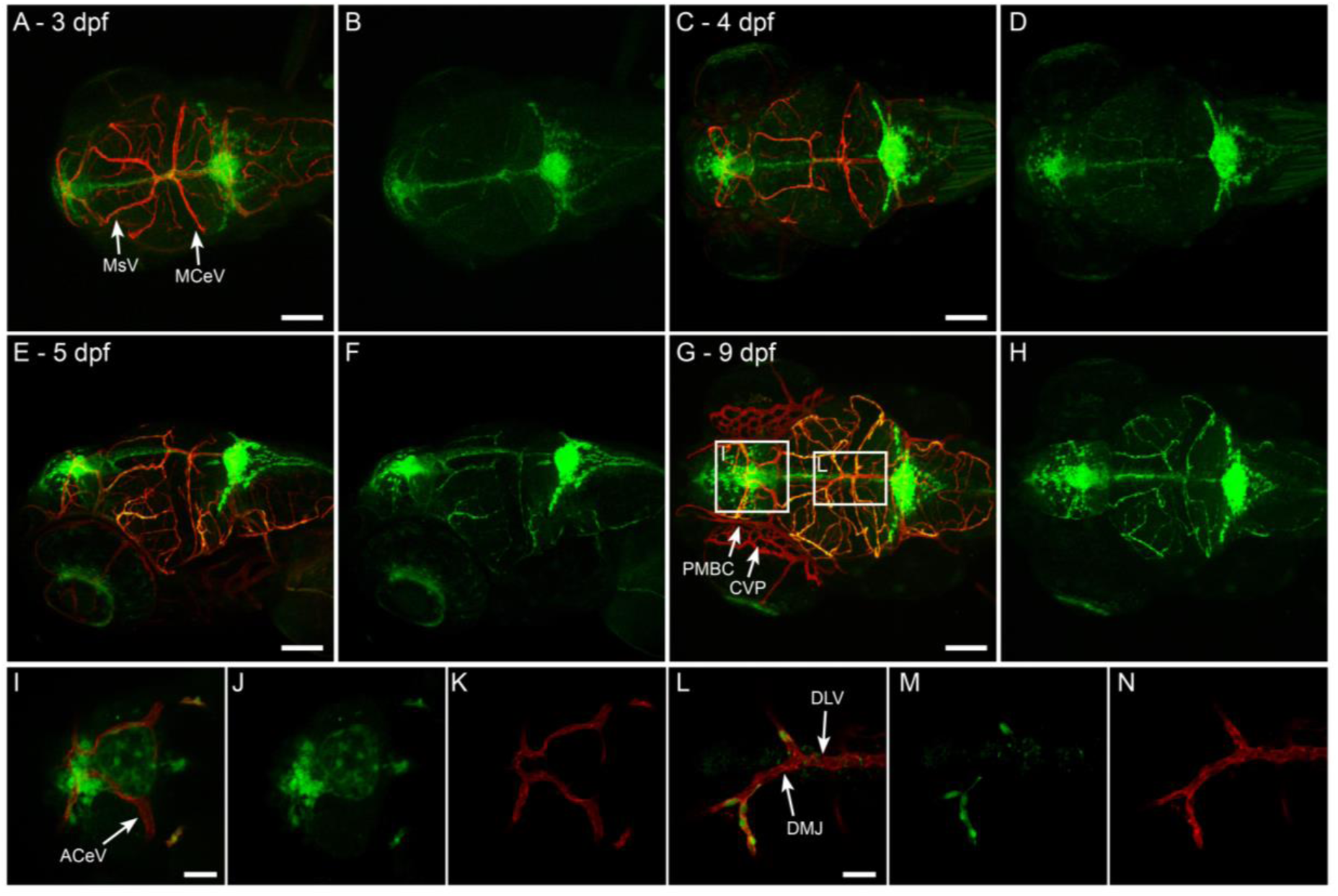
Development of Claudin 5a expression in brain vasculature. A, B. Dorsal view of head of *Tg(kdrl:mCherry)*^*is5*^*;TgBAC(cldn5a:EGFP)*^*vum2*^ larva at 3 dpf. Merge of both channels is shown in A, single green fluorescent expression in B. *Claudin5a* is firstly expressed in the mesencephalic vein (MsV) and middle cerebral vein (MCeV). C-F show expansion of *claudin5a* expression over time, with almost all blood vessels expressing *claudin5a* at 5 dpf (E-F). G,H. Dorsal view of head of *Tg(kdrl:mCherry)*^*is5*^*;TgBAC(cldn5a:EGFP)*^*vum2*^ larva at 9 dpf. Boxed areas are enlarged in I-K and L-N. Specific spots that never express *claudin5a* are: G, closed arrow: primordial midbrain channels (PMBC), choroidal vascular plexus (CVP), I, arrow: anterior cerebral vein (ACeV), L, arrow: dorsal midline junction (DMJ), dorsal longitudinal vein (DLV). Scale bar A-H = 100 μm, scale bar I, L = 25 μm.

### Claudin 5a expression occurs prior to new BBB vessel formation

As stated above, we had identified that Claudin 5a was first present in the larger vessels. Development and expansion of brain vasculature continues after 4 dpf, when *claudin 5a* expression is established. The timing of tight junction protein expression in the BBB has previously been unstudied due to the absence of a suitable *in vivo* model (Haddad-Tóvolli et al., 2017). Thus, we wanted to test our model in this respect and sought to address the question of the timing of tight junction protein expression during angiogenesis of the BBB. Using long time lapse imaging over 12 hours, starting at 96 hpi, in our *Tg(kdrl:mCherry)*^*is5*^*;TgBAC(cldn5a:EGFP)*^*vum2*^ double transgenic we were able to identify sprouting vessels and follow their growth and correlation with *claudin 5a* expression. We hypothesised that *claudin 5a* expression would develop following sprouting/new vessel formation, as we had found in embryonic development (Figure 2). However, careful analysis of sprouting vessels revealed that, in every case, *claudin 5a* expression was observed simultaneously with the initiation of a new vessel (Figure 4, Supplemental Movies S1, and S2). This indicates that components of tight junctions were present from the initiation of BBB angiogenesis and demonstrates the essential nature of early formation of these junctions in BBB function.

**Figure 4:**
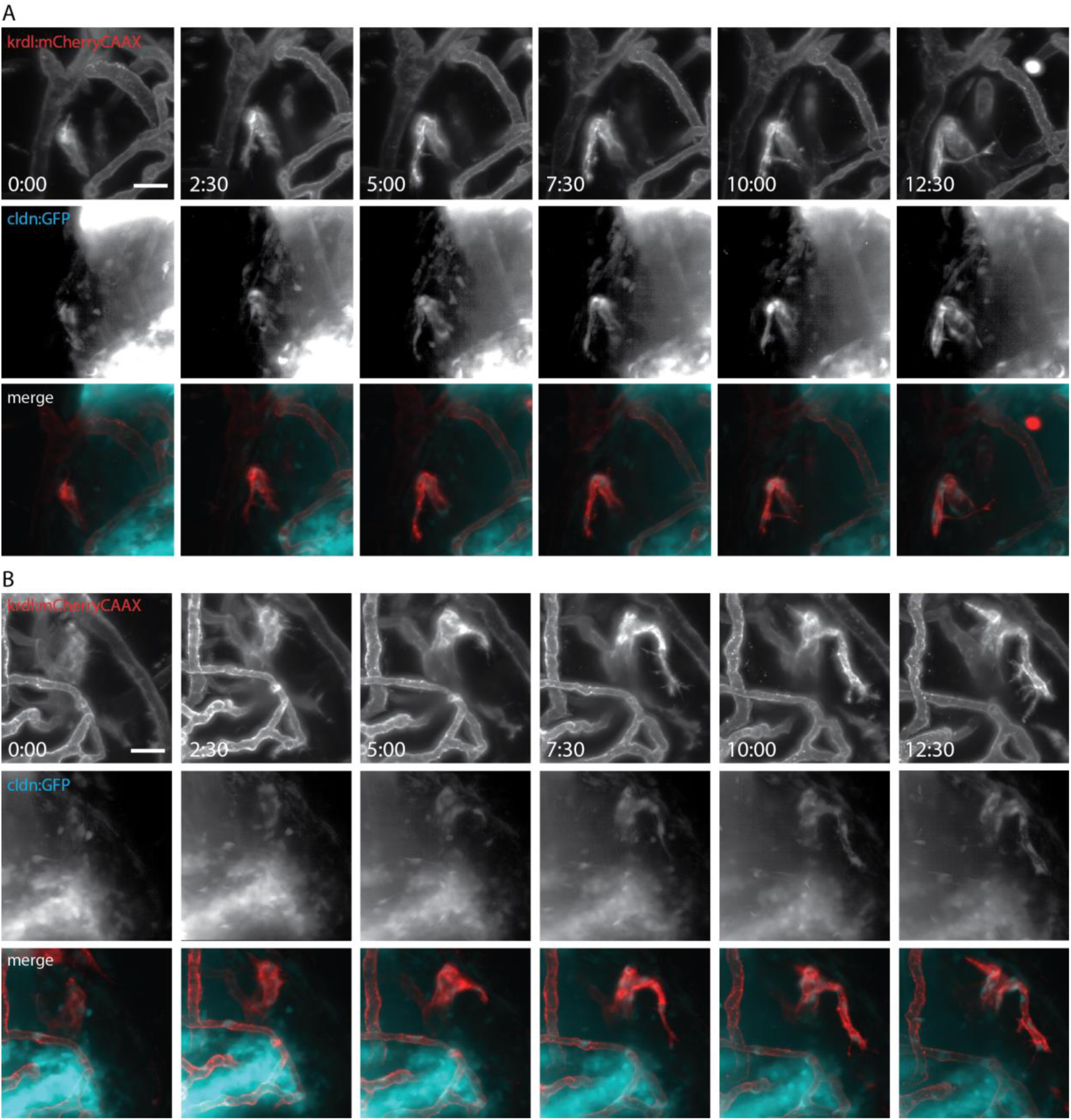
Initiation of *claudin5a* expression co-incides with blood vessel formation. A, B. Still images of 2 time-lapses of blood vessel development in *Tg(kdrl:mCherry)*^*is5*^*;TgBAC(cldn5a:EGFP)*^*vum2*^ larva at 96 hpi. Images were made every 2.5 hours for a period of 12.5 hours. Images of single channels show blood vessel development (*kdrl:mcherryCAAX*) on the top row, co-inciding development of claudin 5a in the middle row and merge image on the bottom row. Scale bar = 20 μm. Corresponding movies of time-lapse can be found in Movie S1 and Movie S2

### Zebrafish larvae possess two separate blood-CP barriers that exhibit collective cell migration

To determine the position of the *claudin 5a* expressing cells being a major component of the blood-CP barrier, its localisation in respect to the vasculature was analysed in more detail. Three-dimensional confocal analysis revealed that the ACeV and prosencephalic artery (PrA) formed a vascular circuit early in development, which overlapped with *claudin 5a* expression at the location of the dCP (Figure 5A, B). In the mCP in the roof of the hindbrain ventricle a similar pattern was seen: the DLV and both posterior cerebral veins (PCeV) formed a vascular circuit closely related to the cells expressing *claudin 5a* (Figure 5C, D). This co-localization did not change during the course of days (Figure 5 A-D, compare 4 dpf with 9 dpf), with the barrier between systemic circulation and CP established early in development. Detailed analysis of the cell dynamics of both CPs identified that the main morphological transformations occur between 1 dpf and 3 dpf (Figure 5E-I), as proposed by a previous study performed in a CP enhancer trap zebrafish transgenic (Bill et al., 2008; García-Lecea et al., 2008). We performed cell-tracking experiments on both CPs, to identify the mechanism of their formation. Cell tracking demonstrated that both structures formed via collective cell migration (Figure 5E, F, Supplemental Movies S3, and S4), where *claudin 5a* expressing cells formed a single layer of closely connected cells that were localized in the roof of both ventricles, before and after movement to the midline (Figure 5 G-I).

**Figure 5:**
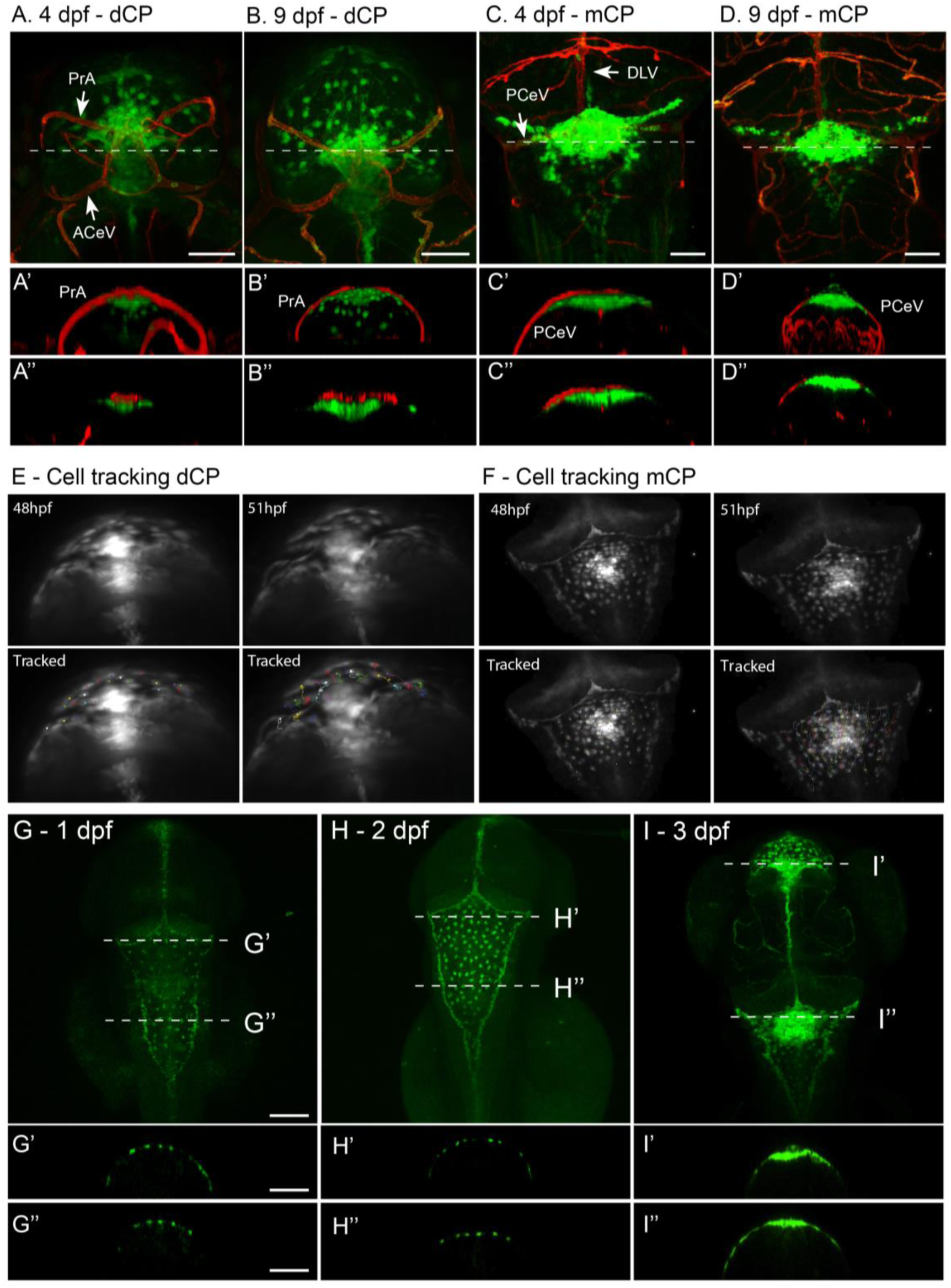
Blood-CP barrier. A. Dorsal view of dCP of larva at 4 dpf, showing the close correlation between the vasculature (red) and *claudin5a* expressing cells (green). The ACeV and PrA form a vascular circuit surrounding the dCP. Transversal view with 3D model (A’) and single Z-slice at the dotted line in A (A’’) visualize the close correlation between these structures. B. Dorsal view of and dCP in a larva at 9 dpf, with transversal view in a 3D model (B’) and single Z-slice at dotted line (B’’). C. At 9 dpf the blood-dCP barrier is formed by the PrA. At the level of the mCP, the PCeV and DLV form a vascular circuit. Transversal view in C’ with a 3D model and C’’ with a single Z-slice at the level of the dotted line in C show the close correlation between the mCP and vasculature. D. Dorsal view of mCP in a larva at 9 dpf, with transversal view in a 3D model (D’) and single Z-slice at dotted line (D’’). Abbreviations: DLV = dorsal longitudinal vein, PCeV = posterior cerebral vein, ACeV = anterior cerebral vein, PrA = prosencephalic artery. Scale bar = 50 μm E. First and last images of time lapse of diencephalic choroid plexus cell migration at 2-3 dpf. Time lapse is presented in Movie S3. F. First and last images of time lapse of myelenchephalic choroid plexus cell migration at 2-3 dpf. Time lapse is presented in Movie S4. G-I. Dorsal view of head of *TgBAC(cldn5a:EGFP)*^*vum2*^ larvae between 1 dpf and 3 dpf, the timeframe in which the major morphological transformation is observed. Transversal section is shown for every time point in panes G’, H’, I’ and G’’, H’’, I’’ corresponding to the dotted lines depicted in G-I. This visualizes the superficial localisation of the GFP expressing cells of the mCP and dCP. Scale bar G = 100 μm

### *Claudin 5a* expression delineates the structured epithelial sheet of the choroid plexus

The choroid plexus is a contiguous epithelial sheet with tight junctions (Lun et al., 2015). To further validate our transgenic, and demonstrate its utility in studying the fine structure of the choroid plexus, we labelled endogenous protein via immunohistochemistry with a monoclonal antibody to mammalian claudin 5 for comparison. Antibody labelling identified a tight network of epithelial cells with claudin 5 localized to the cell margins in both the mCP and dCP structures (Figure 6 A-E). This correlated with *cldn5a*:EGFP expression in the *TgBAC(cldn5a:EGFP)*^*vum1*^ and light sheet imaging was able to resolve the same network of cells and the cell junctions could be identified by a cell thickening at these sites and the resulting increase in EGFP expression (Figure 6 F-J).

**Figure 6:**
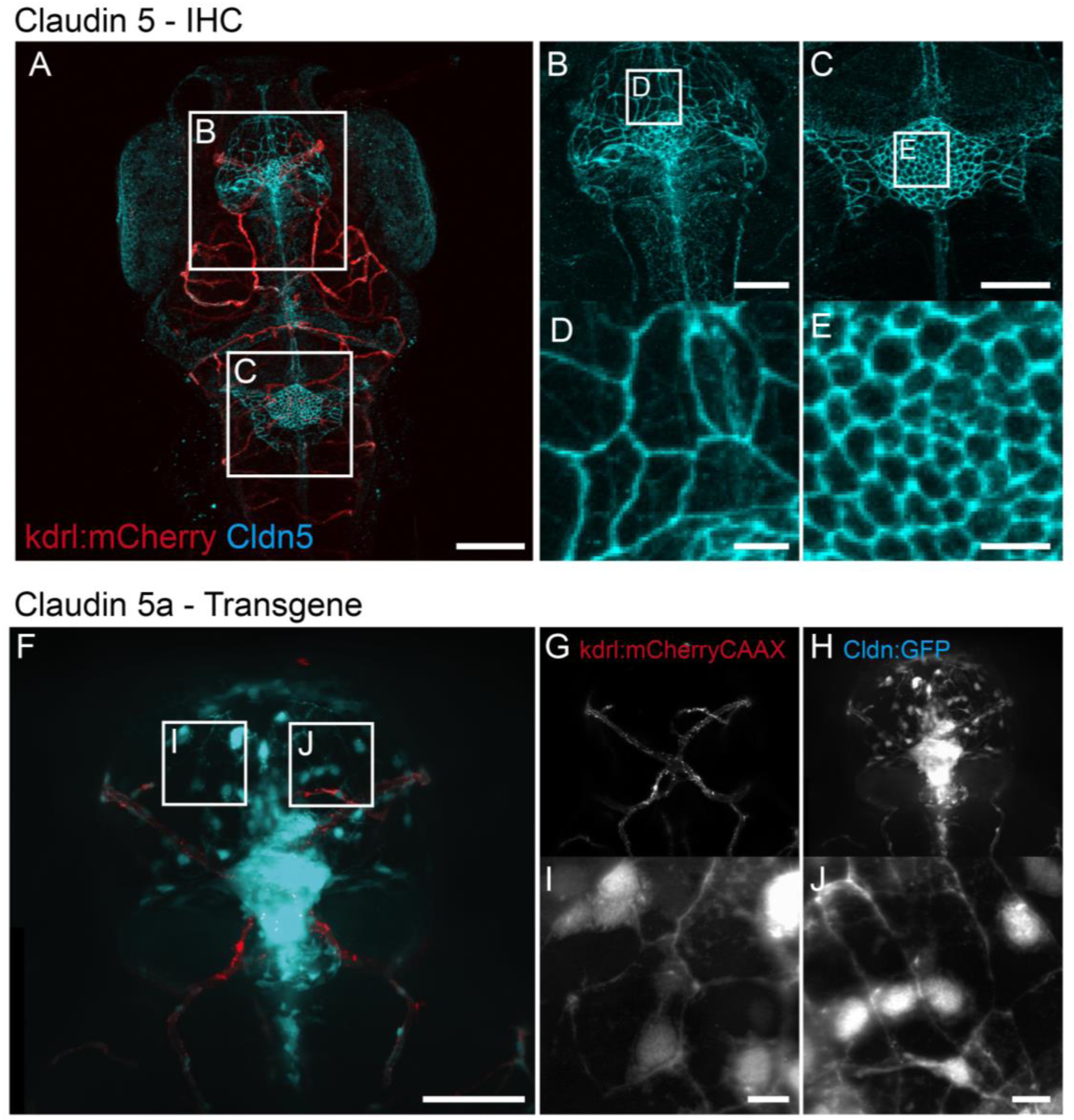
CP morphology – IHC & line. A. Dorsal view of the head of 7 dpf *Tg(kdrl:mCherry)*^*is5*^ larva showing blood vessels (red) and immunohistochemistry labelling of Claudin 5 (cyan). Boxed areas are enlarged in B, D and C, E and show detailed analysis of the network of cells that form the dCP (B) and mCP (C) and is closely connected by Claudin 5. Scale bar A = 100 μm, Scale bar B, C = 50 μm, Scale bar D, E = 10 μm F-J. The network found in *TgBAC(cldn5a:EGFP)*^*vum2*^ larvae is highly similar to the phenotype in A-E. F. Z-stack of mCP of *Tg(kdrl:mCherry)*^*is5*^*;TgBAC(cldn5a:EGFP)*^*vum2*^ larvae with red fluorescent blood vessels (G) and cyan fluorescent *claudin5a* (H). Boxed areas in F are depicted in I and J, which show high magnification of the network formed by *claudin5a* expressing cells forming the mCP. Scale bar F-H = 50 μm, Scale bar I, J = 10 μm

### The *cldn5a*:EGFP expressing epithelial sheet contains ciliated ependymal cells that drive cerebral spinal fluid flow

The cells of the choroid plexus are a specialized type of ependymal cells, which line the brain ventricles. To confirm the identity of our *cldn5a*:EGFP cells we stained for glutamylated tubulin to label cilia. We could image single cilia from *cldn5a*:EGFP cells in the mCP and dCP as early as 2 dpf (Fig 7A-D). Only monociliated cells were found at all stages examined in the fore and hind brain (Fig. 7A-P). We could determine the polarity of the *cldn5a*:EGFP cells on the basis of the abundant labelling of glutamylated tubulin in the skin, which revealed that cilia project into the brain ventricles (Figure 7 B, D, F, H, J, L, N, and P). CSF is under constant flow that is thought to result from a combination of secretion of CSF from the cells of choroid plexus and the beating of the cilia lining the brain ventricles (Kramer-Zucker et al., 2005; Sawamoto et al., 2006). Using injection of fluorescently labelled beads we were able to observe vigorous fluid flow in the CSF in both the fore and hind brain ventricles (Supplemental Movies S5, and S6).

**Figure 7:**
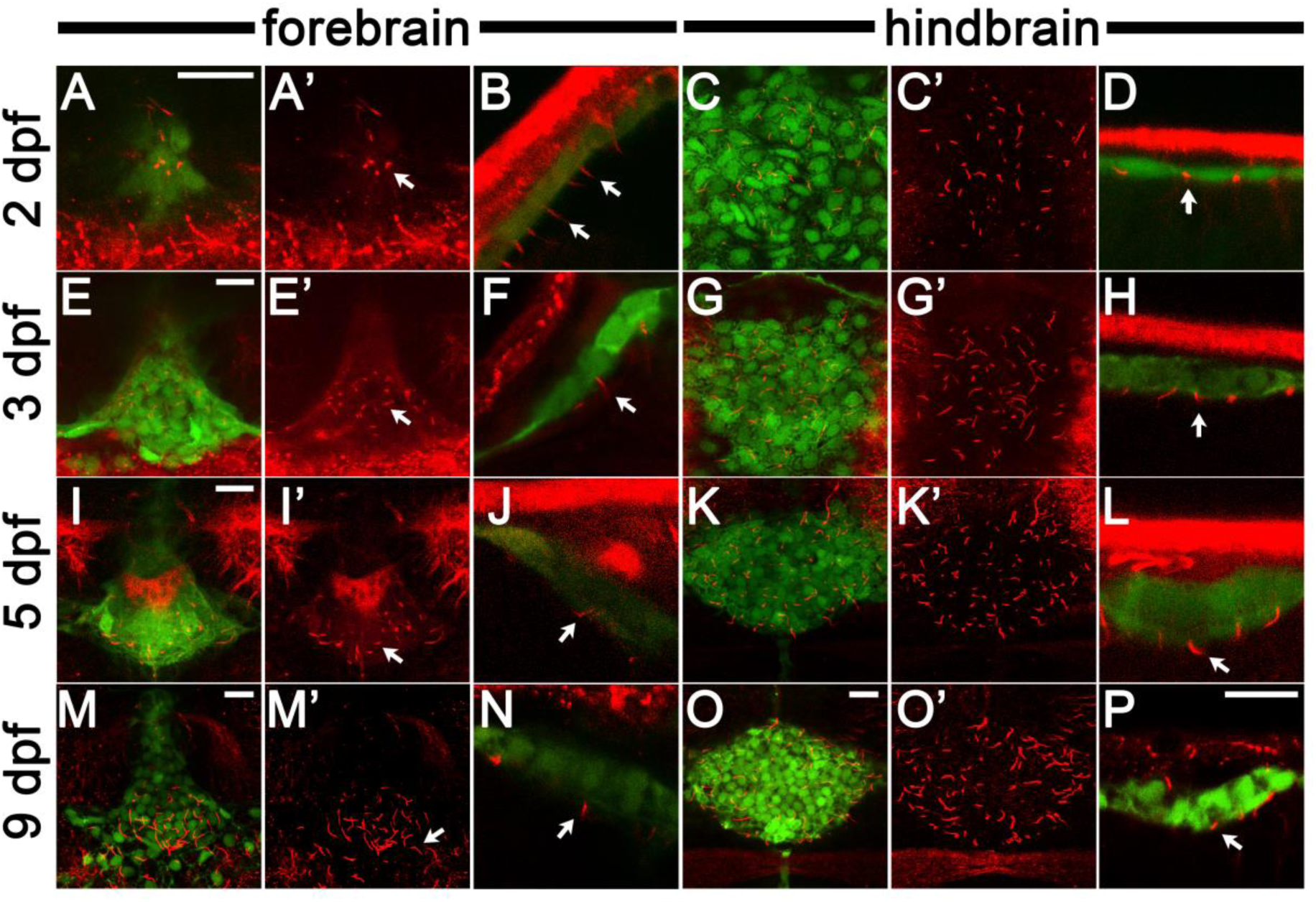
*cldn5a*:EGFP labelled cells in the choroid plexus are ciliated throughout development. Images of cilia visualized by staining with anti-glutamylated tubulin antibody (in red) and *Cldn5a:EGFP* positive cells (in green). (A, E, I, M) Confocal images of the dorsal view of the diencephalic choroid plexus in 2, 3, 5, 9 dpf as indicated. (B, F, J, N) Side views of the diencephalic choroid plexus at the same stages. Cilia project from GFP-positive cells. (C, G, K, O) Dorsal views of the myelencephalic choroid plexus at 2, 3, 5, 9 dpf respectively. (D, H, L, P) Side views of the myelencephalic choroid plexus. Cilia projected from the GFP-labelled myelencephalic choroid plexus cells. For A, C, E, G, I, K, M, O, red channel is shown separately to the right. Arrows indicate cilia. In dorsal view panels, anterior is down. In all side view panels, dorsal is up. Scale bars are 15 micrometres. Scale bar in P applies to all side view images (B, D, F, H, J, L, N, P). Scale bar in O applies to all dorsal view of the myelencephalic choroid plexus (C, G, K, O).

## DISCUSSION

Claudin 5 as prominent TJ protein is a consistent feature between the BBB and blood-CSF barrier (Bill and Korzh, 2014). Here we have used this feature to create an *in vivo* model for real-time analysis of the development, structure and function of the BBB and CP by generating a transgenic zebrafish line that expresses EGFP under the *claudin 5a* promoter. The high homology and synteny with human, the conservation along the teleost lineage and the previous characterisation of Claudin 5 in zebrafish makes *cldn5a* a logical candidate (Abdelilah-Seyfried, 2010; Xie et al., 2010; Zhang et al., 2012).

We show that developmental expression of *cldn5a*:EGFP is restricted to, and starts in both CPs and the midline at 1 dpf, thereby narrowing down the previously shown whole-mount in situ hybridizations (Zhang et al., 2010). The presence of Claudin 5a at the CPs at 1 dpf coincides with the inflation of the ventricles (Zhang et al., 2012, 2010) and corroborates its role in this process. Claudin 5a is besides for tightening the neuroepithelial paracellular barrier probably also important for proper formation of the CP allowing the production of Cerebral Spinal Fluid (CSF) and inflation of both ventricles. These ventricles are connected and form a system through which continues flow of CSF is ensured (Turner et al., 2012). Expression of Claudin 5 possibly outlines the entire ventricular system, which can be an explanation of the midline staining we observe. Expression in the brain vasculature is only found at 3 dpf.

Within the functional highly diverse CNS, the microvasculature is expected to consist of a heterogeneous population of brain microvascular endothelial cells (BMECs) (Wilhelm et al., 2016). A considerable majority of CNS microvasculature comprise capillaries, of which the BMECs preferentially express genes related to transport of ions and nutrients (Macdonald et al., 2010). BMECs of venules instead show higher expression of genes involved in inflammatory-related processes and were shown to have a looser organisation of tight junctions as compared to capillaries. This suggests a vessel specific unique role in physiology and pathophysiology (Macdonald et al., 2010). It is likely that the majority of expression of tight junction related genes cover all vessel types to sustain the protective function of the BBB. Therefore, the observation made in this study that some blood vessel segments lack *claudin5a* expression was highly surprising. In mice, similar heterogeneous expression of claudin-5 has been observed in the spinal cord, with highest expression in capillaries and small venules and less expression in larger venules (Paul et al., 2013). Induction of experimental auto-immune encephalitis (EAE) led to loss of *claudin5* expression specifically in venules, suggesting an important vessel specific role for Claudin 5 in this condition (Paul et al., 2013). Another plausible explanation for the variation in Claudin 5a presence in our model is the anatomical localisation of the blood vessels in respect to brain tissue. Blood vessel segments lacking *claudin5a* expression were all located at the borders of the brain and in close proximity to meninges. Therefore, it is likely that these vessels are located outside the parenchyma and do not possess a BBB.

Development of the CNS vascular network involves complex changes in endothelium and surrounding tissue and the timing of BBB formation in this process is difficult to pinpoint (Malinovskaya et al., 2016). Elaborate studies in rodents have shown that CNS vascularisation during development mainly occurs through angiogenesis derived from the perineural vascular plexus driven by VEGF and CNS-specific Wnt/beta-catenin signalling (Blanchette and Daneman, 2015; Hagan and Ben-Zvi, 2015; Obermeier et al., 2013). Within a few days after initiation of vessel formation, restricted properties have been demonstrated by exclusion of fluorescent dyes from the CNS. Remarkably, this seem to happen before astrocyte generation and ensheathment of vessels occur, while these events have always been considered to be essential for BBB establishment (Blanchette and Daneman, 2015). Moreover, expression of TJ proteins is present at the initiation of angiogenesis in the CNS of mice with subsequent increase of TJ functionality during embryogenesis (Daneman et al., 2010, 2009). In the opossum, it has been demonstrated that newly formed blood vessels possess functional properties from their initiation (Ek et al., 2006). Our study offers a possible mechanism for this, whereby new vessels express *claudin 5a* immediately to form TJs, reflecting developmental steps observed in other models and demonstrating how our transgenic will enable determination of TJ and BBB specification and functionality.

The timespan between initial TJ expression and a functionally intact BBB was for long believed to be the main reason for differences in BBB permeability at different ages. However, considering that these events coincide, an alternative explanation is a prolonged permeability of the barrier between blood and CP in mice (Ek et al., 2006; Saunders et al., 2012). The last decade it has become clear that the blood-CP barrier has more functions than solely CSF production and makes significant contributions to brain homeostasis. Junctional, enzymatic and transporter proteins have been identified and the CP may serve as entry route for immune cells, compounds and even pathogens (Lun et al., 2015). The blood-CP barrier is formed by a monolayer of cuboidal epithelial cells, i.e. ependymal cells, which surround stromal tissue and capillaries, and are joined together by tight junctions (Lun et al., 2015). Studies with enhancer trap lines were the first to describe the two CPs in zebrafish and suggested that at least four different cell lineages develop into stromal, epithelial, endothelial and astroglial components (Bill and Korzh, 2014). The previously described enhancer trap line shows a broad expression profile, of an unidentified gene, in the region of the CP (Bill et al., 2008; García-Lecea et al., 2008). In precision, our transgenic line corroborates and extends those findings and shows cells originating from the roofplate express *claudin 5a* and develop into ciliated ependymal cells.

The treatment of CNS disease is severely impeded by an inability to modulate the entry and exit of therapeutic compounds. Therefore, an improved understanding of the biology of the BBB and blood-CP barrier is of essential importance to improve treatment. Claudin 5 is a tempting target for manipulation, since many hydrophilic drugs prefer to cross the BBB via the paracellular pathway (Greene and Campbell, 2016; Saunders et al., 2013) However, to reduce potential dangerous consequences extensive studies in *in vivo* models are needed before this therapy can be applied in the clinical setting (O’Keeffe and Campbell, 2016). Promising results were achieved in a mice model for cerebral oedema, a major cause for morbidity and mortality in a wide variety of conditions, like severe traumatic brain injury, neurological cancers and brain infections. Strategies to prevent or treat this condition are limited, but transient modulation of claudin-5 with RNAi led to reduced brain swelling and a better outcome (Campbell et al., 2012). In zebrafish, a mutated fragment of clostridium perfringens enterotoxin has been shown to specifically target and modulate claudin 5, inducing transient paracellular permeability of the BBB (Liao et al., 2016). As zebrafish have also proven to be an outstanding model for new compound screens (North et al., 2007; Robertson et al., 2014), analysis of pharmacodynamics of the compound entry into the CNS is now eminently feasible.

## MATERIALS & METHOD

### Identification of Zebrafish Claudin5 and BAC recombineering

Human (genome build GRCh38.p7) CLDN5 protein sequence (NP_003268.2) was used in a BLAST search of the zebrafish protein database (genome build GRCz10). Zebrafish Claudin 5a and Claudin 5b proteins were identified and aligned with human Claudin 5 using Clustal Omega. Using synteny and expression pattern zebrafish *claudin 5a* was confirmed as the homologue of human *cldn5*. A search of Zebrafish genome BACs identified BAC 187M8 from the CHORI211 library (Robert Geisler and Pieter de Jong, Children’s Hospital Oakland Research Institute) as suitable for generation of a fluorescent reporter line due to significant flanking sequence up- and downstream of the *claudin5a* gene. Primers were designed with a forward primer with 50 bp upstream and including the ATG codon of *claudin5a* and 24bp of the targeting vector containing eGFP and a Kanamycin resistance cassette (Dee et al., 2016). The reverse primer contained the reverse complement sequence of the 50 bp downstream of the ATG codon of *claudin5a* and the reverse complement of the end of the cassette sequence. Forward primer: AACTTCTAAACTCCTTTTAGTACCATCAGGAGTGGGAAAAGAAAGCGATGGTGAGCAAGGGCGAGGAGCTGTTC; Reverse primer: GTCCCGCAGACGCACAGGATCAGACCCAGGAGCTCCAAAGCCGCGGAGGCGATATCTGCAGAATTCGCCCTTGA. Tol2 homology arms (Figure S1) were added as described previously (Gray et al., 2011). 2nl of recombineered BAC DNA at a concentration of 50ng/μl combined with itol2 mRNA (Abe et al., 2011) at 30ng/μl.

### Zebrafish

Maintenance of adult zebrafish took place at 26°C in aerated 5 litre tanks, in a 10:14 hour light:dark cycle. Eggs were collected within the first hour post-fertilization (hpf) and injected at the 1-4 cellular stage. Injection was performed as described previously (Benard et al., 2012). Initial transgenesis is performed on: 1) WT zebrafish (ref); 2) casper zebrafish, transparent because these zebrafish lack pigment (White et al., 2008); 3) *Tg(kdrl:mCherry)*, with red fluorescent endothelial cells (Jin et al., 2005). All procedures involving zebrafish embryos and larvae were performed in compliance with local animal welfare laws.

### Transgenesis

At 4 days post-fertilization (dpf) larvae injected with the construct were analysed for transgenic expression with a Leica MZ16FA fluorescence microscope. F0-embryos expressing EGFP in the brain region were selected and grown until reproducing age. Subsequent selection took place and F1 larvae with good expression were used for egg production. F2 larvae were used for further analysis and experiments described here. Stable germline transgenics *TgBAC(cldn5a:eGFP)*^*vum1*^ and *Tg(kdrl:mCherry)*^*is5*^*;TgBAC(cldn5a:eGFP)*^*vum2*^ were generated and used for experiments described in this paper.

### Whole-mount zebrafish larval staining

Visualisation of Claudin 5 expression in the BBB of zebrafish larvae was done by performing whole mount immunohistochemical staining on fixed larvae. For this, larvae were euthanized at indicated time points with tricaine (E10521, Sigma-Aldrich) and fixed in 4% (V/V) paraformaldehyde/PBS (EMS, 100122) at 4°C overnight or at room temperature (RT) for 4 hours in microfuge tubes. Fixed larvae were dehydrated and stored in 100% methanol at −20°C until anti-claudin 5 staining was performed. In short, larvae were rehydrated, rinsed with 1% PBTx (PBS + 1% Triton X-100), permeated in 0.24% trypsin in PBS and blocked for 3 hours in block buffer (10% normal goat serum (NGS) in 1% PBTx (V/V)) Incubation with the primary antibody was done overnight at RT (mouse anti-Claudin 5 (4C3C2), Invitrogen 187364, 1:500 dilution) in antibody buffer (PBTx containing 1% (V/V) NGS and 1% (W/V) BSA). After washing again with PBTx and incubation for 1 hour in block buffer, embryos were incubated in the secondary antibody (goat anti-mouse Alexa-647, Invitrogen A21070, 1:400 dilution), overnight at 4°C. Embryos were then washed with PBTx 5 times, 10 minutes each.

For staining with anti-glutamylated tubulin, washing with PBST (PBS + 0.1% Tween) was applied after fixation. Samples were transferred to 1% Triton X-100 in PBST and incubated at RT for 1-5 days to permeabilise. Larvae were placed in the blocking buffer (0.5% (V/V) Triton, 2% (V/V) normal goat serum in PBST) at room temperature for 2 hours. Subsequently, the blocking buffer was removed and replaced with blocking buffer containing the primary antibody (anti-glutamylated tubulin (GT335) mouse IgG, 1:650 dilution). Specimens were incubated in this solution at 4°C overnight. Larvae were then washed in PBST 3 times, 30 minutes each, on a rotator and incubated in secondary antibody (goat anti-mouse Alexa-568, 1:500 dilution) in the blocking buffer for 4 hours at room temperature. Embryos were washed with PBST 3 times, 10 minutes each.

### Microscopy

For imaging, embryos were mounted in a drop of 1.5% low melting agarose placed on the surface of 1% agarose gel layer in a 35 mm petri dish (Sigma, cat no. A9414) or embedded in 1% low melting-point agarose (Boehringer Mannheim, 12841221-01) dissolved in egg water (60 μg/ml instant ocean see salts) in an 8-well microscopy μ-slide (http://www.ibidi.com). Analysis was performed with a confocal laser scanning microscope (confocal: Leica TCS SP8 X, microscope: Leica DMI 6000). LAS software and ImageJ software were used to generate 3D models, adjust brightness and contrast and create overlays.

### Time lapse imaging

Time lapse imaging was performed with light sheet fluorescence microscopy on a Zeiss Z1 with Z.1 detection optics 20x 1.0NA water immersion objective lens. Zebrafish larvae were mounted in 0.8% low melting point agarose (Cat No. A9414) in E3 containing 0.168 mg/mL tricaine (E10521, Sigma-Aldrich). Z-stacks were captured every 30 minutes over 12.5 or 16 hours. Max intensity projections were generated in Zeiss Zen software. Image processing (cropping, generation of merged images, and linear adjustment of pixel levels) was done in FIJI ImageJ2.0.0. Tracking was performed using the Manual Tracking plugin included in FIJI.

### Bead flow assay

Nacre fish were injected with 4x10^3^ 1.75 mm beads (fluoresbrite carboxylate; Polysciences Inc.) into the hindbrain ventricle 2 days post fertilisation. Bead flow was imaged 24 hours post infection in the hind and fore brain ventricles with a Nikon Ti-E with a CFI Plan Apochromat λ 20X, N.A.0.75 objective lens, and using Intensilight fluorescent illumination with ET/sputtered series fluorescent filters (Chroma, Bellow Falls, VT, USA). Images were captured with Neo sCMOS, 2560×2160 Format, 16.6mm x 14.0mm Sensor Size, 6.5μm pixel size camera (Andor, Belfast, UK) and NIS-Elements (Nikon, Richmond, UK). Images were processed (cropping, contrast enhancement) using NIS-Elements (Nikon, Richmond, UK).

## AUTHOR CONTRIBUTIONS

L.v.L., R.J.E., K.K.J., A.B., J.M., X.F., S.A.J. and A.v.d.S conceived and designed experiments; L.v.L. and S.A.J. made BAC constructs and generated the transgenic lines. T.V. performed zebrafish husbandry. L.v.L., K.K.J did confocal imaging. R.J.E. and A.B. performed light sheet imaging and time lapsing. L.v.L., S.A.J. and A.v.d.S. wrote the paper with input from co-authors.

## ACKNOWLEDGEMENTS

The authors would like to thank Gunny van den Brink – van Stempvoort and Wim Schouten for technical assistance, Wilbert Bitter for valuable input on experimental design and revising the manuscript. We thank the Bateson Centre aquaria staff for their assistance with zebrafish husbandry.

## FUNDING

This research was partially funded by Stichting de Drie Lichten, awarded to L.v.L. RJE was supported by a British Infection Association postdoctoral fellowship. SAJ and AB were supported by Medical Research Council and Department for International Development Career Development Award Fellowship (MR/J009156/1). SAJ was additionally supported by a Krebs Institute Fellowship, and Medical Research Council Center grant (G0700091). Light sheet microscopy was carried out in the Wolfson Light Microscopy Facility, supported by a BBSRC ALERT14 award for light-sheet microscopy (BB/M012522/1).

## COMPETING INTERESTS

The authors declare that they do not have any competing or financial interests.

